# *De novo* whole-genome assembly in interspecific hybrid table grape, ‘Shine Muscat’

**DOI:** 10.1101/730762

**Authors:** Kenta Shirasawa, Akifumi Azuma, Fumiya Taniguchi, Toshiya Yamamoto, Akihiko Sato, Hideki Hirakawa, Sachiko Isobe

**Author notes:** Correspondance: Kenta Shirasawa.

## Abstract

This study presents the first genome sequence of an interspecific grape hybrid, ‘Shine Muscat’ (*Vitis labruscana* × *V. vinifera*), an elite table grape cultivar bred in Japan. The complexity of the genome structure, arising from the interspecific hybridization, necessitated the use of a sophisticated genome assembly pipeline with short-read genome sequence data. The resultant genome assemblies consisted of two types of sequences: a haplotype-phased sequence of the highly heterozygous genomes and an unphased sequence representing a “haploid” genome. The unphased sequences spanned 490.1 Mb in length, 99.4% of the estimated genome size, with 8,696 scaffold sequences with an N50 length of 13.2 Mb. The phased sequences had 15,650 scaffolds spanning 1.0 Gb with N50 of 4.2 Mb. The two sequences comprised 94.7% and 96.3% of the core eukaryotic genes, indicating that the entire genome of ‘Shine Muscat’ was represented. Examination of genome structures revealed possible genome rearrangements between the genomes of ‘Shine Muscat’ and a *V. vinifera* line. Furthermore, full-length transcriptome sequencing analysis revealed 13,947 gene loci on the ‘Shine Muscat’ genome, from which 26,199 transcript isoforms were transcribed. These genome resources provide new insights that could help cultivation and breeding strategies produce more high-quality table grapes such as ‘Shine Muscat’.

## Introduction

Grape (*Vitis* spp., 2*n*=2*x*=38) is one of the most widely cultivated and valuable horticultural crops in the world. World grape production in 2017 was 75,354 kt (FAOSTAT 2019), with the largest producers being China (13,161 kt), Italy (7,170 kt), USA (6,679 kt), France (5,916 kt), Spain (5,387 kt), and Turkey (4,200 kt). This production is mainly for wine, but a considerable portion is also for table fruit use. The most common species used for the production of wine and table grapes around the world is the European grape (*V. vinifera* L.) (Alleweldt and Possingham 1988; Alleweldt, Spiegel-Roy, and Reisch 1991). *V. vinifera* is adapted to a dry and warm climate during its growing season, and countries with a high production of this species have areas with such a climate. However, grapes of this species are highly susceptible to fungal diseases under humid conditions (Hedrick 1908). For example, the failures of early colonists to establish *V. vinifera* in the United States resulted from a lack of resistance to native diseases, soil pests, and low winter temperatures in the northernmost areas (Einset and Pratt 1975). To overcome this difficulty, seedlings or selections from wild American native grape species that survived under North American conditions were collected and many breeders attempted to improve the American native species through hybridization among *V. vinifera* and American native species in the latter half of the 19^th^ century (Snyder 1937). The species most often used as a cross parent was *V. labrusca* (the fox grape), which provided disease and cold resistance as well as a distinctive flavor (Einset and Pratt 1975).

*V. labruscana* L.H. Bailey is defined as a subgroup of grapes that originated from hybridization of *V. labrusca* with other species, most commonly *V. vinifera* (Bailey and Bailey 1930), and more than 1,500 *V. labruscana* varieties, such as ‘Campbell Early’, ‘Catawba’, ‘Concord’, ‘Delaware’, and ‘Niagara’, have been developed (Hedrick 1908; Snyder 1937). Japan has a humid climate throughout its growing season and Japanese grape crops are susceptible to attack from fungal diseases during periods of rainfall. This climate limitation has hampered the development of *V. vinifera* production in Japan. Consequently, Japanese grape breeders have attempted to develop new cultivars through interspecific hybridizations to combine crisp flesh and favorable flavor traits derived from *V. vinifera* grapes, with ease of cultivation (mainly disease and berry cracking resistance) from *V. labruscana* grapes.

‘Shine Muscat’ is a promising cultivar in Japan. Grape breeding in the Division of Grape and Persimmon Research, Institute of Fruit Tree and Tea Science, National Agriculture and Food Research Organization (NARO), Japan, used a long-term multigenerational strategy to develop ‘Shine Muscat’ as well as other cultivars. An offspring grape with crisp flesh, Akitsu-21 (*V. labruscana* × *V. vinifera*), was obtained by crossing ‘Steuben’ (*V. labruscana*) with ‘Muscat of Alexandria’ (*V. vinifera*). Akitsu-21 was then crossed with ‘Hakunan’ (*V. vinifera*) to develop ‘Shine Muscat’ (*V. labruscana* × *V. vinifera*) (‘Shine Muscat’ grape breeding group in Institute of Fruit Tree and Tea Science 2018; Yamada, Yamane, and Sato 2017; Yamada et al. 2008). Because of its favorable fruit eating quality when produced under Japanese climate and environmental conditions, together with its sophisticated cultivation system (http://www.naro.affrc.go.jp/laboratory/nifts/shine-muscat/cultivation-method.html), the cultivation area of ‘Shine Muscat’ rapidly increased after its release by NARO in 2006 and reached 1,196 ha in 2016 (Ministry of Agriculture, Forestry and Fisheries of Japan 2019). The ‘Shine Muscat’ grape cultivar has large yellow-green berries, crisp flesh texture, muscat flavor, high soluble solids concentration and low acidity. Berries can be eaten with the skin. It is moderately tolerant to downy mildew and ripe rot, but sensitive to anthracnose (Kono et al. 2013; Shiraishi M et al. 2007; Yamada et al. 2008). The fruit shelf life is longer than that of ‘Kyoho’ (*V. labruscana* × *V. vinifera*, 2*n*=4*x*=76), the current leading variety in Japan, and cold-hardiness is comparable with that of ‘Kyoho’.

Owing to the importance of grape in food industry, grapevine (*Vitis vinifera*) was the first bearing fruit, the second tree, and the fourth higher plant for which whole-genome sequencing was reported (Jaillon et al. 2007; Velasco et al. 2007). The original genome project used a *V. vinifera* cultivar, ‘Pinot Noir’. Since the original genome publication, the ‘Pinot Noir’ genome sequence data has been updated and the last version was released in 2017 (Canaguier et al. 2017). In addition, to our knowledge, *de novo* genome sequence assemblies have been released for four wine cultivars of *V. vinifera*: ‘Cabernet Sauvignon’ (Chin et al. 2016; Minio, Massonnet, Figueroa-Balderas, Vondras, et al. 2019; Minio et al. 2017), ‘Carménère’ (Minio, Massonnet, Figueroa-Balderas, Castro, et al. 2019), ‘Chardonnay’ (Zhou et al. 2018; Roach et al. 2018), and ‘Zinfandel’ (Vondras et al. 2019). In parallel, the *Vitis* pan-genome was released, in which whole-genome sequences of as many as 472 accessions of 48 species were sequenced to find structure variations as well as sequence variations across the *Vitis* diversity panel (Liang et al. 2019). However, little genome information for *V. labruscana* × *V. vinifera* hybrids is available at present. This is probably due to the complex nature of the *V. labruscana* × *V. vinifera* genomes owing to their interspecific hybridizations.

Recent advances in sequencing technology and data analysis methods have made it possible to dissect the complexity of genomes of interspecific hybrids as well as polyploids (Kyriakidou et al. 2018). There are two strategies for sequencing complex genomes. One is to determine “unphased” sequences, in which a consensus sequence is obtained from the two or more genomes comprising the hybrids/polyploids by collapsing the homologous/homoeologous sequences or choosing one of the redundant sequences. The other approach is “phased” genome sequencing to determine all haplotype/subgenome sequences of the multiple genomes in hybrids/polyploids. Haplotype phasing is therefore an essential component of phased sequencing analysis. Long read sequencing technologies provide phase information as single reads are of sufficient length to cover more than two sequence variant sites and allow haplotype determination. A genome assembler, FALCON-Unzip (Chin et al. 2016), for PacBio long reads based on single molecule real-time sequencing determines two types of genome sequences from the hybrids/diploids: primary contigs, contiguous sequences representing the haploid genome; and haplotigs, fragmented sequences generated from heterozygous genome regions. Furthermore, FALCON-Phase extends the contiguity of haplotigs using contact information provided by Hi-C data (Kronenberg et al. 2019). Triobinning, an alternative assembly strategy to that of FALCON, is able to resolve two haplotypes prior to assembly (Koren et al. 2018). These long-read-based sequencing approaches are effective for characterizing the complex genomes of plants (Jiao and Schneeberger 2017; Li and Harkess 2018). Short-read technologies can also be employed for analysis of complex genomes. The linked-reads provided by Chromium technology (10x Genomics) generate a “pseudo” long read by assembling short reads from a long DNA molecule. In an F1 hybrid of pepper, haplotype-phased genome sequences were constructed by assembling the “pseudo” long reads (Hulse-Kemp et al. 2018). Another short-read-based method is DenovoMAGIC (NRGene). Whereas details for the DenovoMAGIC algorithm have not been described, the contiguity of the resultant sequences offers substantial improvements over other technologies in terms of the numbers of plants sequenced, including crops with hybrid/polyploid genomes (International Wheat Genome Sequencing et al. 2018; Edger et al. 2019; Hu et al. 2019; Chen et al. 2019). More recently, an open-source short-read assembly tool, TRITEX, was developed for polyploid genomes (Monat et al. 2019).

Genomics has the potential to enhance cultivation and breeding strategies toward production of high-quality fruits, vegetables, and flowers with attractive consumer-friendly phenotypes. With the advanced technologies and methods available, it becomes possible to achieve sequencing analysis of the complex interspecific hybrid genomes often observed in elite cultivars of horticultural crops such as the ‘Shine Muscat’ table grape. In this study, we determined both “phased” and “unphased” genome sequences of ‘Shine Muscat’, an interspecific grape hybrid (*V. labruscana* × *V. vinifera*). In addition, we used full-length transcriptome sequencing to gain insights into a gene set associated with elite table grape characteristics.

## Materials and methods

### Plant materials

For genome assembly, DNA was extracted from young leaves (expanded to ∼6 cm) collected from a vine of ‘Shine Muscat’ (*V. labruscana* × *V. vinifera*) growing in a vineyard at the Division of Grape and Persimmon Research, Institute of Fruit Tree and Tea Science, NARO, Higashihiroshima, Hiroshima, Japan. For transcriptome analysis, total RNA was extracted from young leaves, tendrils, flower clusters before flowering, and skins, flesh, and seeds of mature berries at harvest.

### Genome sequence analysis

Paired-end (PE) libraries (insert sizes of 450–470 bp and 700–800bp) and mate-pair (MP) libraries (insert sizes of 2–4 kb, 5–7 kb, and 8–10 kb) were constructed with the KAPA Hyper Prep Kit (Kapa Biosystems, Roche) and Nextera Mate Pair Library Preparation Kit, respectively. A 10x Genomics Chromium library was also prepared. The libraries were sequenced on a HiSeq 2500 system (Illumina) in PE, 250-bp mode (for the PE library, insert sizes of 450–470 bp) and a NovaSeq 6000 system (Illumina) in PE, 150-bp mode (for the remaining five libraries). The PE reads were used for genome size estimation based on k-mer frequency with Jellyfish (Marcais and Kingsford 2011). Sequencing data were processed and assembled using DeNovoMAGIC3 to generate both unphased and phased genome sequences. Small sequences not integrated into the assemblies were gathered as unplaced sequences. The integrity of the assemblies were verified using the Benchmarking Universal Single-Copy Orthologs (BUSCO) method (Simao et al. 2015). Comparative analysis of the genome structures was performed using D-GENEIS (Cabanettes and Klopp 2018).

### Transcriptome analysis

Total RNAs extracted from six tissues of ‘Shine Muscat’ were mixed to prepare an Iso-Seq library in accordance with the manufacturer’s protocol (PacBio). The library was sequenced with single molecule real-time sequencing technology on a Sequel system. The obtained reads were clustered using the Iso-Seq 3 pipeline implemented in SMRT Link v. 6.0 (PacBio), mapped on the unphased sequence of the ‘Shine Muscat’ genome with Minmap2 (Li 2018), and collapsed to obtain nonredundant isoform sequences using a module in Cupcake ToFU (https://github.com/Magdoll/cDNA_Cupcake). Functional annotation was assigned to the isoforms with Hayai-Annotation (Ghelfi et al. 2019).

## Results

### Sequencing and assembly

DNA from leaves of ‘Shine Muscat’ was sequenced. In total, 120.4 Gb and 106.6 Gb sequence data were obtained from Illumina PE and MP libraries, respectively, as well as 34.0 Gb from a 10x Genomics Chromium library. The distribution of distinct *k*-mers (*k*□= 17) showed two peaks at multiplicities of 111 and 225, indicating heterozygous and homozygous regions, respectively (Figure 1). This suggested that the heterozygosity of the ‘Shine Muscat’ genome was high, indicative of an interspecific hybrid. Haploid (n) size of the ‘Shine Muscat’ genome was estimated to be 493.0 Mb and diploid (2n) size was estimated to be 999.2 Mb (Figure 1).

**Figure 1.**
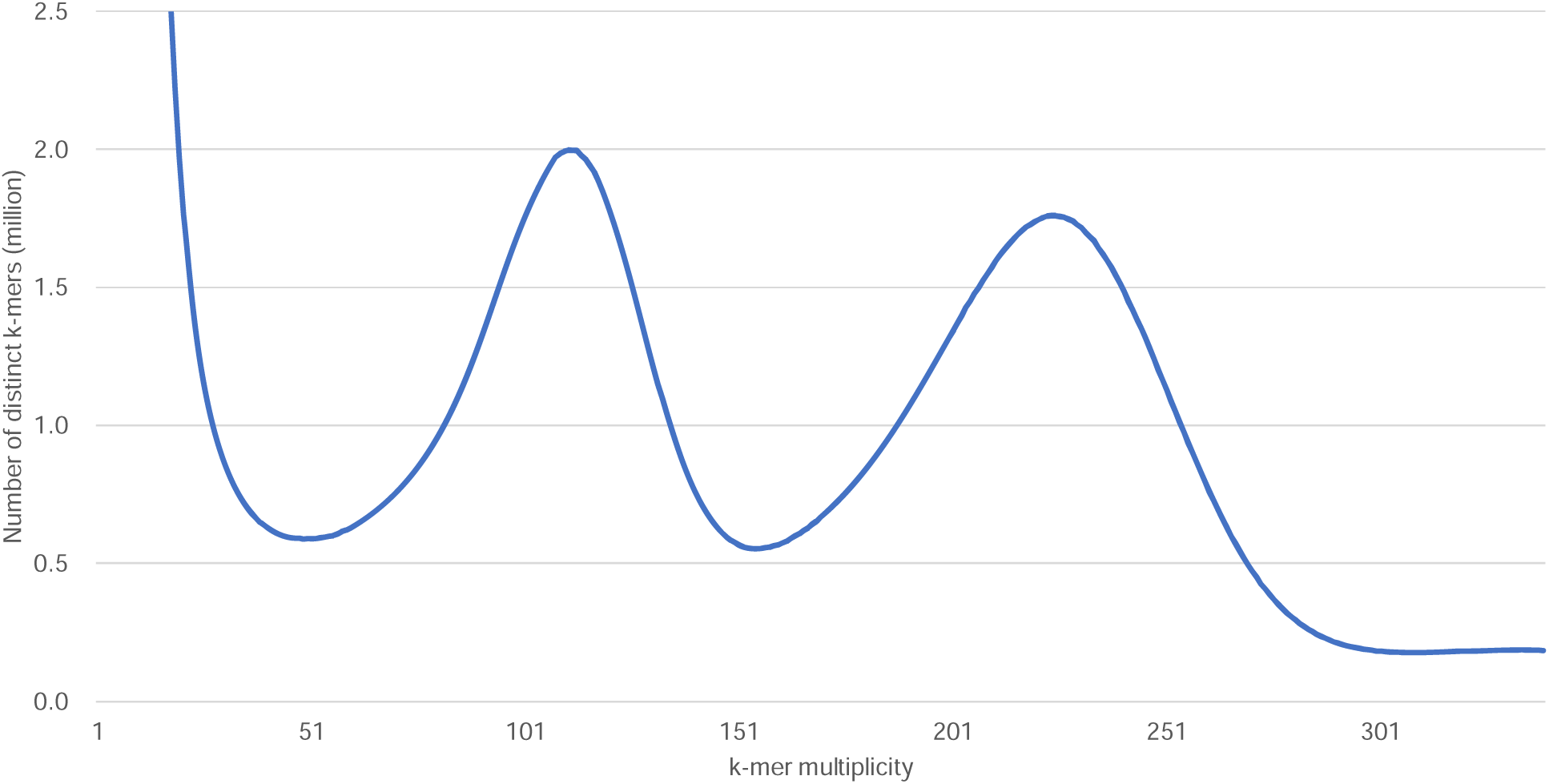
Genome size estimation for ‘Shine Muscat’ with the distribution of the number of distinct *k*-mer (*k*=17) with the given multiplicity values.

The sequence reads from the PE, MP, and 10x Genomics Chromium libraries were assembled to obtain unphased and phased genome sequences, designated VSMuph_r1.0 and VSMph_r1.0, respectively (Table 1). VSMuph_r1.0 consisted of 8,696 scaffold sequences covering 490.1 Mb with N50 of 13.2 Mb and N90 of 2.5 Mb. The assembly included 94.7% complete BUSCOs (92.5% single copy and 2.2% duplicated). VSMph_r1.0 was shorter than VSMuph_r1.0 and had 15,650 scaffolds spanning 1.0 Gb with N50 of 4.2 Mb and N90 of 214 kb. VSMph_r1.0 contained 96.3% complete BUSCOs (21.2% single copy and 75.1% duplicated). As expected, single-copy and duplicated BUSCOs were predominant in VSMuph_r1.0 and VSMph_r1.0, respectively. The remaining unassembled 361,173 contig sequences, spanning 175 Mb, were collected as VSMupl_r1.0.

**Table 1.**
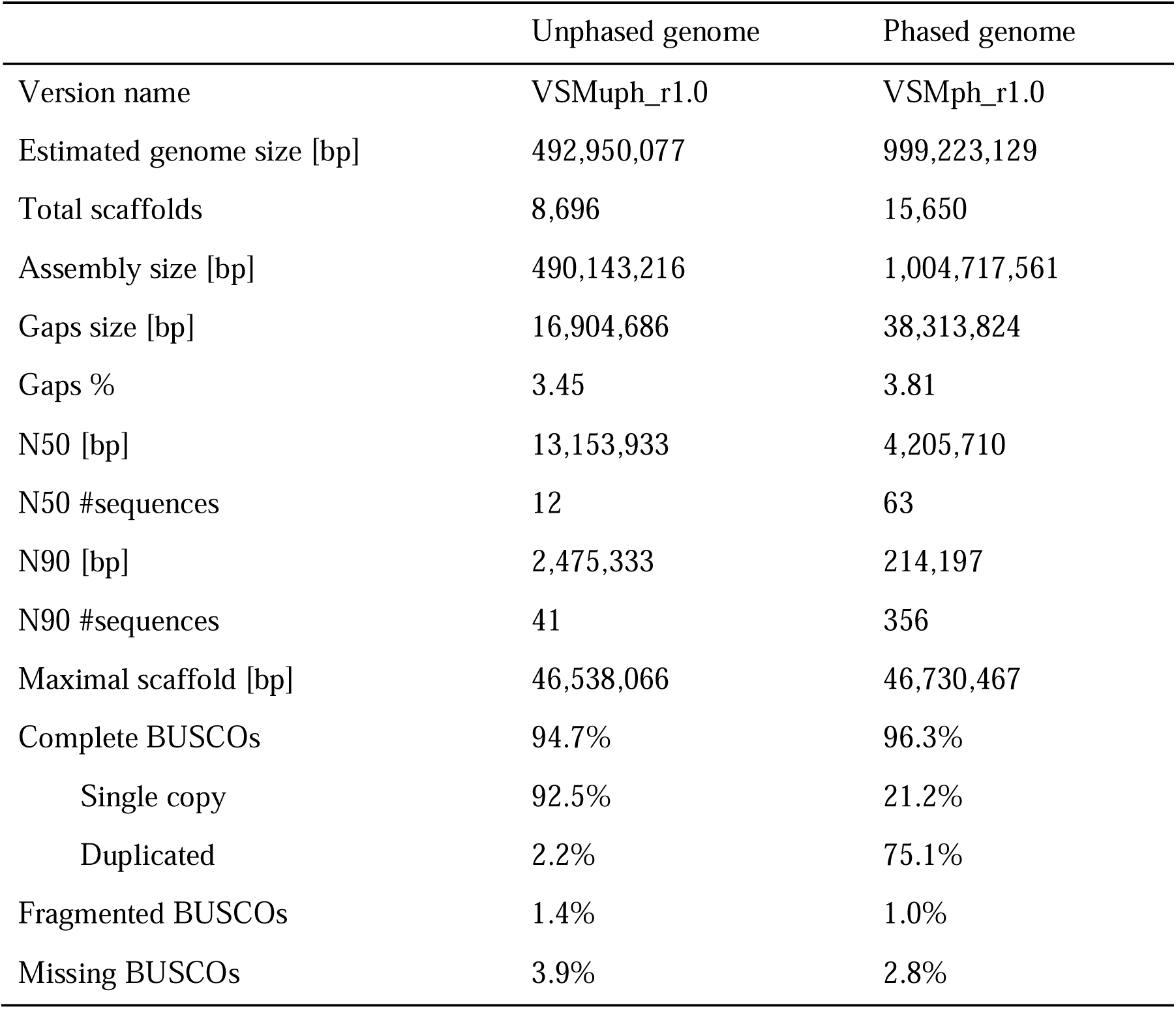
Assembly statistics of the ‘Shine Muscat’ genome sequences.

The VSMuph_r1.0 assembly covered the entire VSMph_r1.0 genome, and vice versa, indicating that both sequences represented the same region of the ‘Shine Muscat’ genome (Figure 2A). As expected, the phased VSMph_r1.0 assembly was approximately double the size of the unphased VSMuph_r1.0 assembly. To assess completeness of coverage over the *Vitis* genome, both assembly sequences were compared with the pseudomolecule sequence (chromosome 1 to 19) of *V. vinifera* ‘Pinot Noir’ (version 12X) (Canaguier et al. 2017). This analysis confirmed that VSMuph_r1.0 and VSMph_r1.0 completely encompassed the reference sequence of the *V. vinifera* genome (Figure 2B and 2C), and confirmed the doubled size of VSMph_r1.0 (Figure 2C). However, in parallel, no collinear relationship was observed in chromosomes 1, 2, 4, 6, 7, 9, 10, 11, 12, 14, 15, 17, 18, and 19 for VSMuph_r1.0 (Figure 2B) and chromosomes 6, 7, 9, 15, 17, 18, and 19 for VSMph_r1.0 (Figure 2C), suggesting potential structure variations between the genomes of the two species genomes or possible technical errors during assembly of the ‘Shine Muscat’ and/or ‘Pinot Noir’ genome sequences.

**Figure 2.**
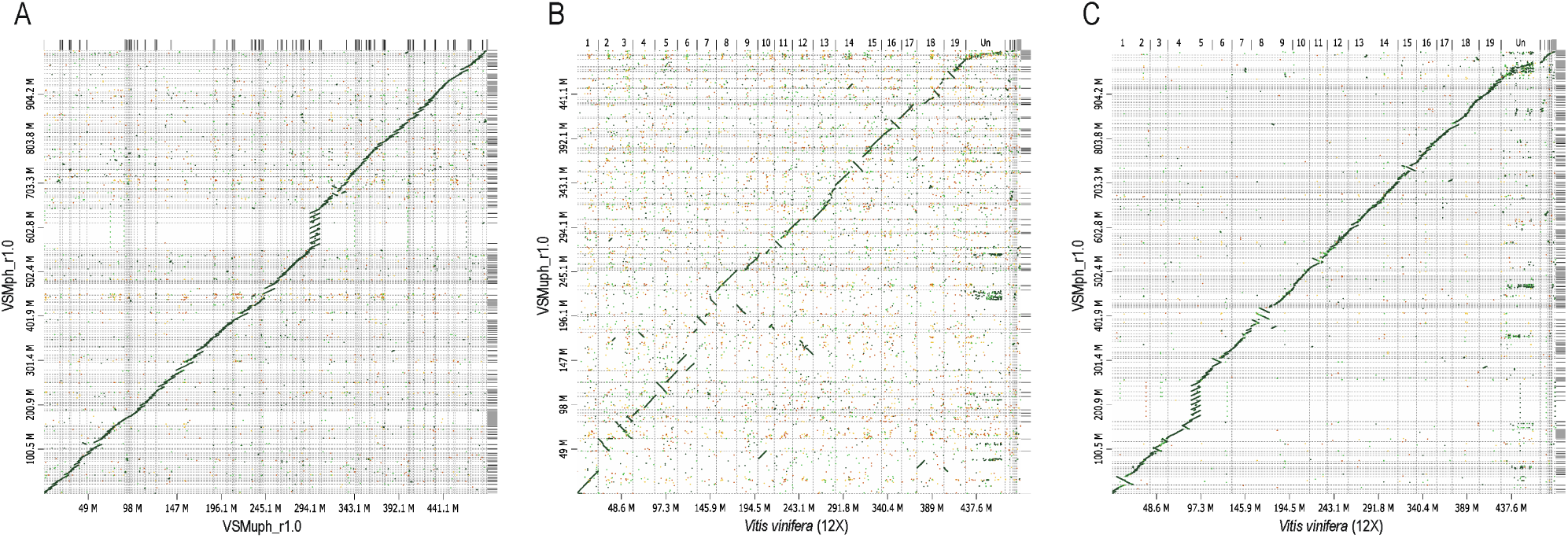
Synteny relationships of the grape genomes. Syntenic regions are indicated with plots between the unphased and phased genome sequences of ‘Shine Muscat’ (A) and between the ‘Pinot Noir’ genome, version 12X and either the unphased (B) or phased (C) ‘Shine Muscat’ genomes.

### Gene discovery through full-length transcriptome sequencing

RNA was sequenced from six ‘Shine Muscat’ tissues: young leaves, tendrils, flower clusters before flowering, and skins, flesh, and seeds of mature berries at harvest. In total, 68.8 Gb sequence data consisting of 57,425,099 sequence reads were obtained. The reads were clustered using the ISO-Seq3 pipeline to obtain 68,709 high-quality isoforms. The isoforms were then mapped onto the unphased VSMuph_r1.0 genome and redundancy was collapsed. In total, 26,199 isoforms were identified at 13,947 gene loci (1.9 isoforms per gene locus) across the unphased genome, of which 72.0% complete BUSCOs (53.7% single copy and 18.3% duplicated) were represented. Of the 26,199 gene isoforms, 11,659 (44.5%), 13,896 (53.0%), and 15,577 (59.5%) were assigned to Gene Ontology Slim terms in the biological process, cellular component, and molecular function categories, respectively, while 2,269 isoforms had enzyme commission numbers (Supplementary Table S1).

## Discussion

Here, we report the first draft genome sequence of an interspecific hybrid table grape, *V. labruscana* × *V. vinifera*. ‘Shine Muscat’, an elite table grape cultivar bred at NARO and cultivated across Japan (‘Shine Muscat’ grape breeding group in Institute of Fruit Tree and Tea Science 2018; Yamada, Yamane, and Sato 2017; Yamada et al. 2008), was chosen for sequence analysis due to its high heterozygosity resulting from interspecific hybridization (Figure 1). Haplotype-phased genome sequences representing the two progenitor genomes were determined as well as the “unphased” reference genome sequence. *V. vinifera* was one of the earliest plants to be sequenced, and due to technical limitations at that time, initial genome sequencing focused on inbred diploid lines (*V. vinifera* PN40024) (Canaguier et al. 2017; Jaillon et al. 2007). Subsequent advances in sequencing technologies allowed in-depth analysis of heterozygous genomes in some wine grape cultivars (Chin et al. 2016; Minio, Massonnet, Figueroa-Balderas, Castro, et al. 2019; Roach et al. 2018; Vondras et al. 2019; Zhou et al. 2018). However, genome sequences for table grapes, which are often derived from interspecific hybrids, have not been available to date. In this study, advanced techniques were used to successfully sequence the genome of an interspecific table grape hybrid. Sequence contiguity and quality, which were supported by N50 length and BUSCO analysis, respectively (Table 1), compared favorably with those of the reported grapevine genomes (Canaguier et al. 2017; Chin et al. 2016; Jaillon et al. 2007; Minio, Massonnet, Figueroa-Balderas, Castro, et al. 2019; Roach et al. 2018; Velasco et al. 2007; Vondras et al. 2019; Zhou et al. 2018).

Potential structure variants were identified between the genomes of ‘Shine Muscat’ and ‘PN40024’ (Figure 2). These structure variants could be responsible for phenotype differentiations between the two cultivars, or two species, including the characteristics that influence crop use. To verify and support this hypothesis, more experimental data would be required. To this end, we have started genetic mapping and Hi-C analysis for ‘Shine Muscat’ to construct two chromosome-level pseudomolecule sequences representing complete haplotype-phased genomes. Further transcriptome analysis to complement the current data (Table 2) is underway to allow identification of genes with high accuracy and coverage. Alongside a pan-genome approach (Liang et al. 2019), the ‘Shine Muscat’ sequence data will enhance our understanding of the composition and structure of the ‘Shine Muscat’ progenitor sequences. This resource may allow future identification of genes or genetic loci that contribute to the phenotypic differences between the two cultivars, or between table- and wine grapes. Thus, the availability of the whole-genome sequence of the interspecific hybrid *V. labruscana* × *V. vinifera* will considerably facilitate future molecular genomic and genetic research in this area. The ‘Shine Muscat’ genome will act as a platform upon which future genetic studies can build, allowing selection of favorable traits from interspecific hybrid grapes and contributing to the future development of novel interspecific hybrid varieties.

This first draft genome sequence of the interspecific hybrid *V. labruscana* × *V. vinifera* will also facilitate studies into the physiology and biochemistry of berry development and maturation in table grapes. Ongoing genomic and transcriptomic analysis will further improve the utility of these genetic resources and provide new insights into cultivation and breeding strategies for high-quality table grapes like ‘Shine Muscat’.

## Availability

The sequence reads are available from the DNA Data Bank of Japan (DDBJ) Sequence Read Archive (DRA) under the accession number DRA008743. The DDBJ accession numbers of assembled scaffold sequences, VSMuph_r1.0, are BKBX01000001–BKBX01008696. Sequence similarity searches by BLAST (Altschul et al. 1990) to the genome assemblies and transcripts and a genome browser based on JBrowse (Buels et al. 2016) are available at Plant GARDEN (https://plantgarden.jp).

## Acknowledgments

This research was supported in part by the Kazusa DNA Research Institute Foundation and Cabinet Office, Government of Japan, Cross-ministerial Strategic Innovation Promotion Program (SIP), “Technologies for Smart Bio-industry and Agriculture” (funding agency: Bio-oriented Technology Research Advancement Institution, NARO).

## Supporting information

**Supplementary Table S1** Functional gene annotation for isoforms from Iso-Seq analysis

